# SeqStruct: A New Amino Acid Similarity Matrix Based on Sequence Correlations and Structural Contacts Yields Sequence-Structure Congruence

**DOI:** 10.1101/268904

**Authors:** Kejue Jia, Robert L. Jernigan

## Abstract

Protein sequence matching does not properly account for some well-known features of protein structures: surface residues being more variable than core residues, the high packing densities in globular proteins, and does not yield good matches of sequences of many proteins known to be close structural relatives. There are now abundant protein sequences and structures to enable major improvements to sequence matching. Here, we utilize structural frameworks to mount the observed correlated sequences to identify the most important correlated parts. The rationale is that protein structures provide the important physical framework for improving sequence matching. Combining the sequence and structure data in this way leads to a simple amino acid substitution matrix that can be readily incorporated into any sequence matching. This enables the incorporation of allosteric information into sequence matching and transforms it effectively from a 1-D to a 3-D procedure. The results from testing in over 3,000 sequence matches demonstrate a 37% gain in sequence similarity and a loss of 26% of the gaps when compared with the use of BLOSUM62. And, importantly there are major gains in the specificity of sequence matching across diverse proteins. Specifically, all known cases where protein structures match but sequences do not match well are resolved.

## Introduction

Proteins are the central point players on the field of biology, and a deeper understanding of their behaviors will facilitate the more meaningful interpretation of genome data, particularly for drawing evolutionary conclusions. The present work utilizes the Big Data of protein sequences and structures, which have grown rapidly ^2^, to develop a more reliable way to link between sequences and phenomes. Rapid progress in genome sequencing has already provided hundreds of millions of protein sequences, and similar advances in structural biology now provide over 100,000 protein structures, which are deemed to be a nearly complete set of characteristic structures (folds). Comparative modeling can produce homologous structures for many of the remaining sequences having unknown structures ^3-17^, but still in many cases is unable to identify good template structures because of inadequacies in sequence matching. Sequence matching is extremely important because it is a highly efficient way to compare proteins (genes), identify protein (gene) functions and permits rapid comparisons and analyses of whole genomes.

### What is missing in sequence matching?

Sequence matching is based on the accumulated statistics of amino acids changes in the individual columns of multiple sequence alignments; these data have been used for generating amino acid similarity matrices such as the BLOSUM series of matrices^18^. Globular proteins are packed at high densities inside their structures, an important point made many years ago by Fred Richards ^19^ and others ^20^. The average packing density of amino acids is 0.74, which is about the same as the highest packing density of close packed spheres. The direct implication of this high packing density is that substitutions will be interdependent. Previously we and others showed how the packing density of amino acids is overall related to sequence entropies and consequently to residue conservation ^21, 22^. That is, in general, densely packed amino acids are more conserved. The challenge, however, is to account properly for the complexities of the packing within the amino acid similarity matrix, since these interdependences have not previously been considered and yet can strongly affect the possible substitutions.

Our group has long experience in mining interaction information from protein structures, and we have demonstrated how accounting for 3-body and 4-body interactions can improve the empirical pairwise interactions potentials ^23-25^, because these more adequately represent the correlations within the high-density proteins. These interaction potentials have proven to be successful ways to assess the quality of predicted protein structures at CASP. Here we will perform a similar extraction to learn about which pairs of amino acids in structures are more readily substituted with other pairs of amino acids. Then, we go ahead to show how accounting for these pair exchanges significantly improves nearly all protein sequence alignments.

### Previous amino acid similarity matrices

The first similarity matrices developed were the PAM matrices developed by Margaret Dayhoff ^26^ based on a very small number of sequences. The most widely used at present are the BLOSUM matrices ^18^ developed by Henikoff and Henikoff ^18^. Another type of substitution matrix based on amino acid contact frequencies was reported by Miyazawa and Jernigan ^27^. Muller et al. ^28^ developed the VTML substitution matrices. Others considered particularly the positions that are different in different families of proteins ^29^. Yamada and Tomii ^30^ recently reported a matrix based on principal component analysis and the variabilities across previous substitution matrices. But, none of these approaches account for the interdependences of substitutions, and individually they all yield remarkably similar results.

There is a history of using protein structures to develop sequence alignment matrices, but none of these have been particularly effective, nor have they come into common usage. Since structures are more conserved than sequences ^31^, an appropriate way to approach this problem could be by the use of structure alignments. Prlic *et al.* used structure alignments to derive similarity matrices (PRLA1) ^32^. They used a data set of superimposed protein pairs to derive evolutionary information. These pairs had high structural similarity but low sequence similarity. Structural information has also been used to enrich substitution matrices ^33^ by using a linear combination of the sequence substitution matrix BLOSUM50 and a threading energy table. The resulting matrix was shown to improve prediction accuracy for homology modeling in the twilight zone. The Johnson and Overington matrix (JOHM) took into account not only the substitutions that occur in similar parts of protein structures but also accounts for the variable regions where gaps occur^34^. Blake and Cohen built similarity matrices (*e.g*.: BC0030) where structural superposition of protein structures was performed by using structures obtained from the CATH database ^35^. Structures were selected based on the sequence identities and the alignments were performed for different ranges of sequence identities. These have been used in structure-function predictions. Some other studies have shown that the use of protein-family-specific substitution matrices is helpful to identify orthologs that are not identifiable with the standard BLOSUM matrices ^36^. Recently we also explored this approach, but have still obtained relatively small gains ^37^.

## Method

We take an entirely new unique approach to this problem by deriving a universal amino acid similarity matrix that incorporates the effects of a large set of pairwise substitutions. First we extract sequence pair correlations from the multiple sequence alignments for 2,005 Pfam domains. Then we filter these correlations to retain only the pairs of amino acids closest together (and interacting most strongly) in the structures. We name this new amino acid similarity matrix **SeqStruct** for **S**equence **C**orrelated, **S**tructurally **C**ontacting substitutions. Deriving a simple amino acid similarity matrix based only on this data thus includes the effects of the correlated pairs expressed in terms of their effects on single amino acid matches. This new matrix is then combined additively with the BLOSUM62 matrix to include also the singlet amino acid similarities.

### Extract Sequence Correlations

The large body of sequence data contains the important details about co-evolving residues. Correlated mutations (compensatory mutations) result from the fact that amino acid substitutions at a given position in a dense protein environment can be compensated by other pairs of mutations. In these pairs, there can be either single amino acid changes reflecting the likely exchange of one member of the interacting pair within the context of the second, or changes in both positions reflecting within the particular protein context. For example, at two positions in a structure, a large-small pair of amino acids might be substituted for a small-large pair without disrupting the structure. Such co-evolving pairs are identified from Multiple Sequence Alignments (**MSA**) of sets of thousands of related sequences. These types of correlations have recently proven to be highly useful for predicting interacting pairs of residues for structure predictions ^38-43^. Note that these correlations can lead to substitutions of one type of amino acid by another type that would not usually be considered to be so similar. But, this is where the major strength of this new approach lies - in picking up these non-intuitive contextual changes. A schema for this approach is shown in **Fig. 1**. This leads immediately to a list of ranked pairwise substitutions.

**Figure 1.**
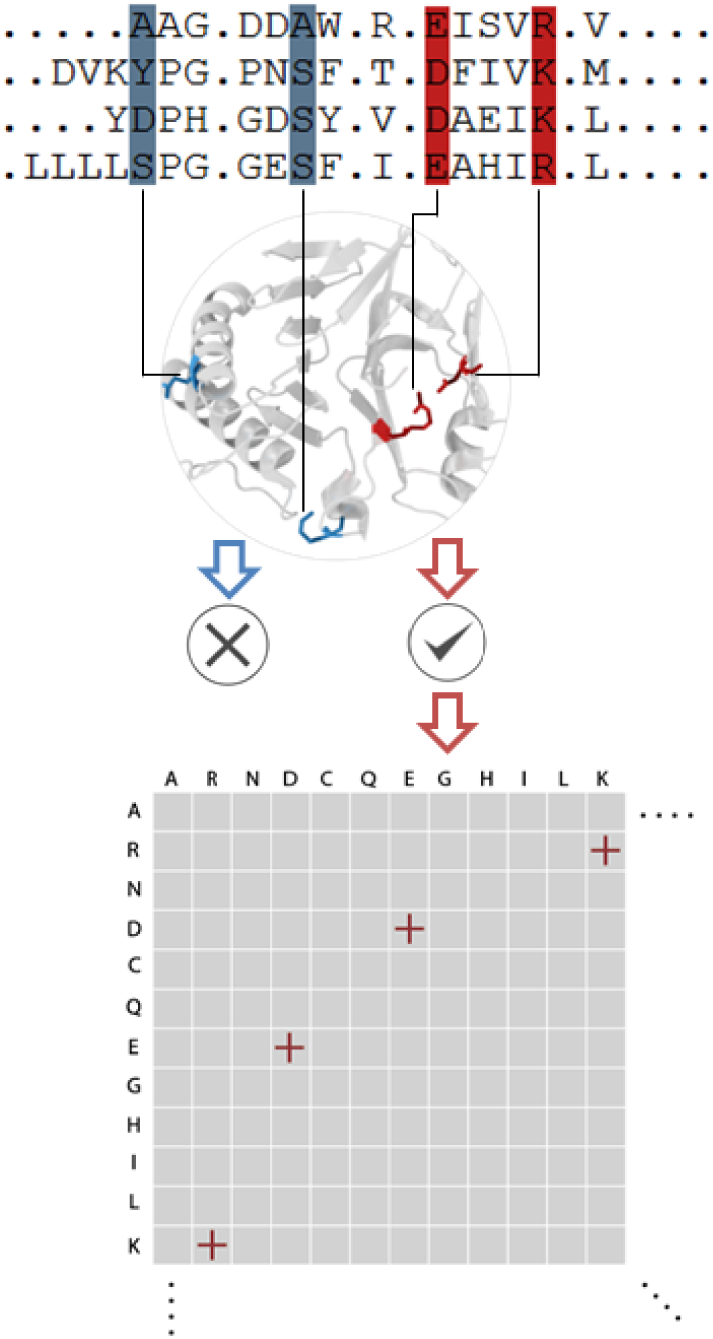
Derivation of the SeqStruct amino acid similarity matrix. Pairs of residue positions having correlations greater than 0.75 in the multiple sequence alignments are filtered by retaining only the correlated pairs in close contact (red) and discarding those not in close proximity (blue). 2,005 Pfam domains were used to derive the sequence correlations. The resulting **SeqStruct** substitution matrix is based on the cumulative information from all of these structural domains (see Supplementary **Table S1** for the list of domains whose MSAs are used, together with the representative structures)

### Retaining only the correlations for proximate interacting pairs

Correlations among sequences can originate from many different factors including long-range allostery and functional considerations specific to a given type of protein. Here we take the expedient measure of limiting the correlations to those for physically adjacent and directly interacting residues in the structures. For this purpose we utilize the atoms on the ends of the amino acid side chains. These atoms have previously been used by the Liang group ^44, 45^ to achieve higher specificities in empirical potential functions and for characterizing binding sites. This represents a coarse-graining of the amino acids at the level of one or a few geometric points per amino acid. We have long experience in using coarse-graining to extract empirical potentials of interactions from protein structures ^24, 27, 46-55^. This coarse-graining is essential for making direct connections between structures and sequences. This is a successful approach, and we have extensive experience with such approaches ^56-58^.

It is clear that such proximate amino acid pairs should have the strongest interactions. The large amount of sequence and structural data means that there is sufficient data even after imposing such a stringent limit on the included pairs. Collecting this data closely resembles what is done for the empirical energy potentials for interacting amino acids, where our research group has extensive experience ^49, 59-65^. Results from CASP have definitively demonstrated that using these “statistical” knowledge-based potentials of the type pioneered by the Jernigan group is by far the most successful approach for evaluating structure predictions ^11, 12, 66-69^. The previous generation of protein potentials suffered in general from not being sufficiently specific, and specifically using larger groups of 3 or 4 interacting residues with direction-dependent potentials derived from a large dataset of protein structures ^55^ has improved these. Presumably further gains will be possible for amino acid similarities by incorporating correlations of groups larger than simple pairs, and the sequence data can provide the huge amount of data required for this.

### Co-evolution detection from multiple sequence alignments

The co-evolution data from multiple sequence (MSA) can be used for different purposes. One of the most successful applications of the MSA data has been for protein structure prediction. Mutual information ^70^ was introduced as a straightforward approach for predicting residue proximity in structures. It is based on the association of information entropy between a pair of residues. The results are confounded by indirect cases correlated with intermediate positions. Methods have been developed to remove such indirect or transitive effects. (Here we are doing something similar, but much simpler, by simply removing the correlated pairs that are not in direct contact in the pdb structure representative of a particular Pfam domain.) Such methods include direct coupling analysis (DCA) ^41, 71^ and protein sparse inverse covariance (PSICOV) ^72^. These both established a global statistical model for using multiple sequence alignments to extract the position-specific variabilities and inter-position correlations. One notable recent study ^73^ has compared these methods. Amino acid co-evolution information derived from MSAs has been used to predict residue proximity; here we use this data instead to identify the correlations in positions in a MSA. Another example of the use of mutual information is the identification of the catalytic amino acids in enzymes ^74^. In that approach, higher order associated amino acids from a group of residues with high pairwise co-evolving couplings were connected by transitivity. Also, the statistical coupling analysis (SCA) ^75, 76^ explicitly searches for groups of co-evolving residues that specify the allosteric pathways within a protein structure.

### Develop Structural Frameworks for the Use of Big Sequence Data

Structures provide a highly specific, and physically certain, framework for mounting these correlated pairs. Instead of focusing only on the pairwise couplings from a statistical amino acid contact prediction method, the results here identify the co-evolving amino acids that specifically interact in the strongest ways in the structures. For each multiple sequence alignment in Pfam ^77^ a representative structure is identified. This structure and its allosteric properties add a physical interpretation to the sequence correlations. Our data is collected to ensure both strong correlations as well as proximity (and strong interactions) within the structures. These correlated pairs modify the individual amino acid substitutions in complex ways, accounting for the effects of pairwise amino acid changes. In this way, we obtain a universal amino acid similarity matrix that is superior to those presently in use.

### Use frequencies as logs of probabilities

For extracting correlations we will follow the well-established approach ^47^ of taking logs of frequencies. The correlations extracted will then have the important property of being additive rather than multiplicative. This is essential for incorporating the correlations into the SeqStruct amino acid similarity matrix developed here. The result is a quantum leap in the capabilities of sequence matches to provide fast and meaningful information about protein function from sequence alone. This will lead to important solutions to many existing problems in the identification of protein (gene) function in biology.

### Substitution matrix derivation

The elements in an amino-acid substitution matrix are log-odds scores, which is the logarithm of the ratio of the likelihoods of two hypotheses: one amino-acid should be replaced by another amino-acid and one amino-acid should not be replaced by another. The log-odds scores describe the tendency of one amino-acid may or may not be substituted by another amino-acid in the homolog protein sequences. Among all the co-evolution based methods, mutual information has been widely applied and is well studied due to its computational simplicity and the explicit in measuring the co-evolution dependence. The mutual information between two positions in a multiple sequence alignment is defined as following:

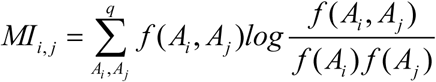

where *q* is the number of alphabets in the multiple sequence alignment; *f* (*Ai*) is the frequencies of an amino-acid *A* has been observed on the position *i*. *f* (*Ai*, *Aj*) is the observed co-occurring frequencies for two amino-acids on position *i* and position *j*. Both single and the co-occurring frequencies are weighted by the similarity of the sequences in the multiple sequence alignment.

Here, we calculate the mutual information upon a large set of multiple sequence alignments which corresponding to 2,005 Pfam protein families. In order to obtain reliable results, each of the selected multiple sequence alignments is chosen to include at least 1,000 sequences. For each protein family, a set of position pairs having significant mutual information are selected to compare against the residue contacts for the corresponding protein structure. The position pairs having significant mutual information AND an observed residue contact are considered to contain both genuine co-evolution dependences and important structural information. The corresponding columns of the multiple sequence alignments are saved for the amino-acid substitution extraction.

To explore a variety of the new substitution matrices, we also adopt the following scheme from ^37^ to derive a set of combined matrices:

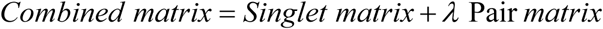

The “Singlet matrix” is taken as BLOSUM62, the most widely used of such matrices. In this way we derive a set of combined substitution matrix that includes both singlet and pairwise information.

In this project, we search different values and found the range of λ from 2 to 5 for the best performance when the Original matrix is taken to be BLOSUM62. Finally, the scores in the matrix are rounded to the nearest integer for more efficient computing. The value of λ = 2.5 yields the resulting best **SeqStruct** matrix shown in **Fig. 2**. λ was evaluated with 22,765 pairs of non-redundant sequences (having less than 90% identity). The values there show major changes that reflect the effects of the correlated substitutions.

**Figure 2.**
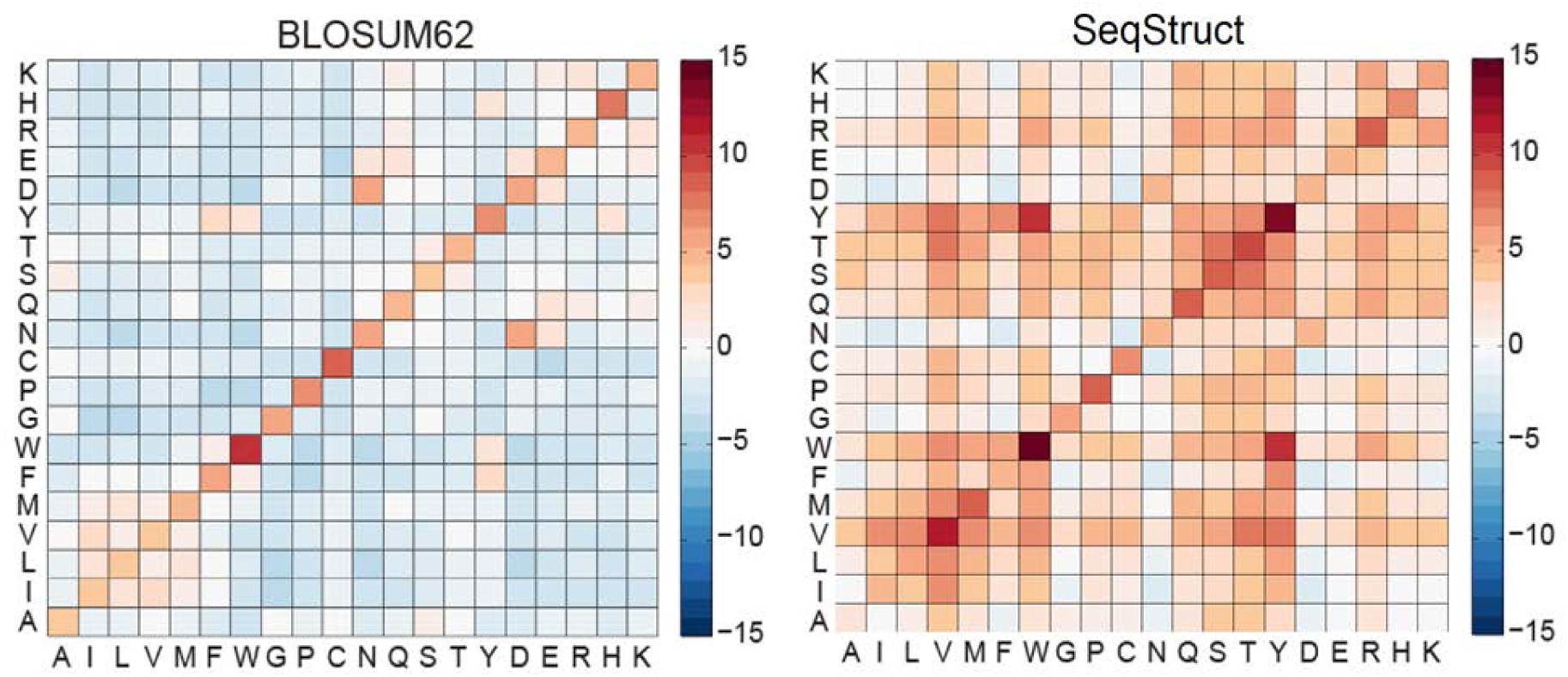
Comparison of the new SeqStruct amino acid similarity matrix with the BLOSUM62 matrix. Red values are more conserved than blue values. There are significant differences introduced into the extent of conservation. W, Y and V are the most conserved residue types, and Y-W interconversions are likewise favorable. There are significantly more substitutions favored (redder values) in the new matrix than in BLOSUM62.

### Congruence to the Statistical Characteristics of a Structural Domain

The best global alignments are obtained with the rigorous Needleman-Wunsch dynamic programming approach ^78^, which we have used throughout. This method begins with an initial scoring matrix for every possible match with this two-dimensional scoring matrix. The method uses dynamic programming to choose the best global match as the best combination of alignments of these pairs. ***Because the new* SeqStruct *amino acid similarity matrix includes pairwise information, the scoring procedure is equivalent to a 3D procedure because it accounts for the important pairs of substitutions within the structures.*** Once a query sequence is scored against different protein families the quantitative scores will immediately identify the best alignment(s).

Gap penalties have been investigated to obtain the best scores, but the results show extremely low sensitivity to these parameters As a result we conclude that there is no need to include gap penalties.

## Results

This new **SeqStruct** amino acid similarity matrix was tested for 3,160 sequence matches (see online paper for details). The dynamic programming Needleman-Wunsch sequence alignment method ^78^ was utilized; results are shown in **Fig. 3**. In almost every case the number of identities and similarities increased over the use of BLOSUM62 and the number of gaps was reduced. But, importantly the sequence similarity showed major gains. A summary of these results is show in **Table 1**, with striking result of an increase of 37% in the similarities and a reduction of gaps by 26%.

**Table 1.**
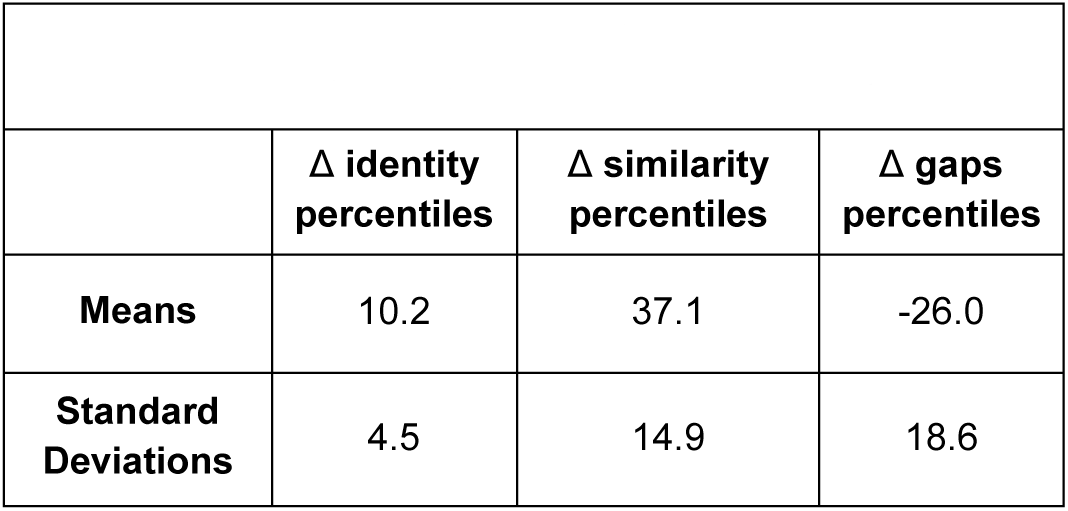
Means and standard deviations of gains in sequence matching for 3,160 matches

**Figure 3.**
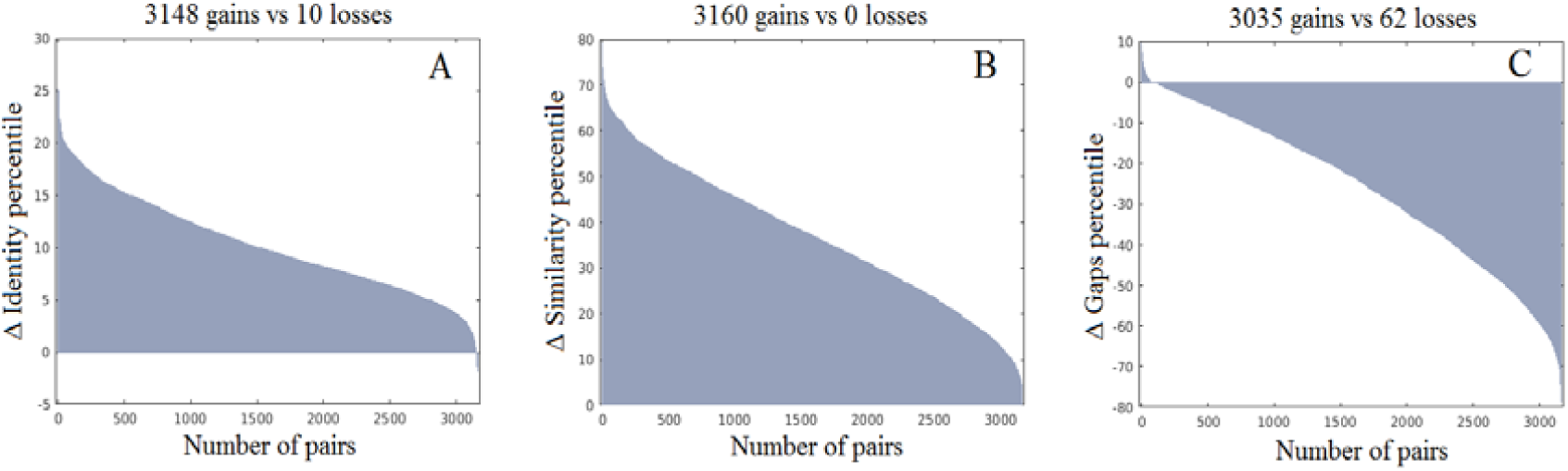
Gains sequence identity and sequence similarity, together with loss in the numbers of gaps in the sequence alignments with SeqStruct over BLOSUM62 for 3,160 sequence matches.

There are a substantial number of known cases where two protein structures are similar but their sequences are not. We have used the set of such protein pairs identified by Friedberg and Margalit ^1^ and remarkably they all are found to be substantially more similar with the use of the new **SeqStruct** amino acid similarity matrix. The results clearly demonstrate that the new **SeqStruct** matrix resolves many of these problematic cases. Examples of these improved alignments is shown in **Fig. 4**, showing also how well the aligned sequence segments are positioned in the structure alignments.

**Figure 4.**
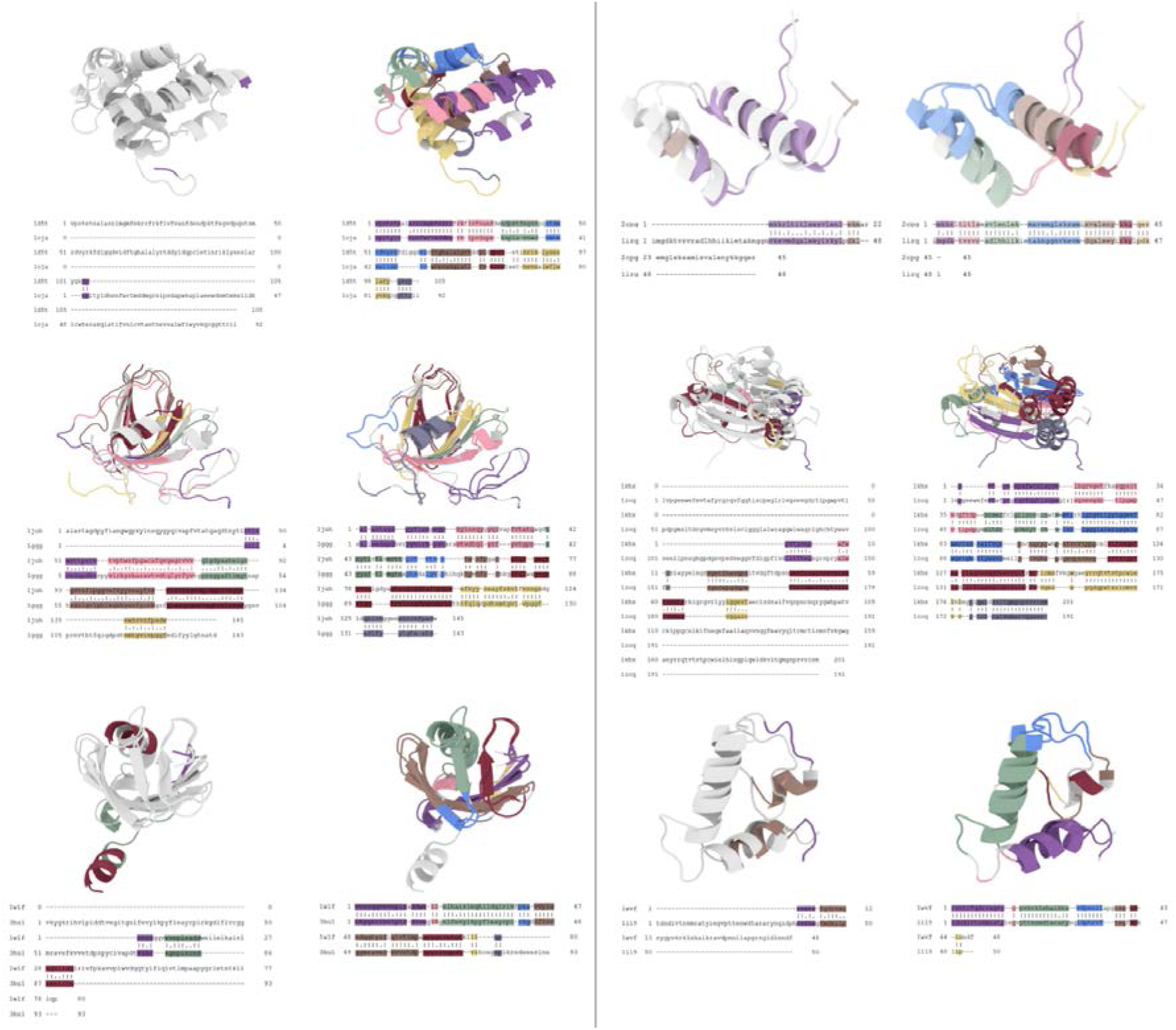
BLOSUM62 fails to identify similar structures, but our new amino acid similarity matrix successfully identifies the specific structure pair. Sequence and structure matches are compared, and structures are colored according to matched segments in the sequence matches. Eight cases are shown. On the left in each case is the alignment performed with BLOSUM62, and on the right the alignment with our new **SeqStruct** amino acid similarity matrix. The segments in gray correspond to the parts aligned with gaps in the sequence alignment. This are taken from the 92 hard cases that were identified in Reference ^1^ together with an additional 174 cases that we have identified.

The results shown **in Figs. 2-4 and Table 1** are based on including the effects of the sequence changes for the individual singlet substitutions, together with pair-wise correlations derived from the MSAs and filtered to be only those sites that are close to one another in the corresponding protein structures. These show major gains over the use of BLOSUM62 alone for several reasons: 1) combining Big Sequence Data with structures, 2) using the Pfam definitions of protein domains and the corresponding MSAs and 3) the way in which the sequence correlations are screened for directly interacting pairs. This improvement to the performance of protein sequence matching derives significantly from incorporating structural information into an amino acid similarity matrix.

## Discussion

While the results have been transformed into a single amino acid substitution matrix having dimensions of 20×20, the available data could be used to explicitly consider substitutions of pairs, thus requiring larger matrices. It remains to be seen whether such a larger computational effort would be worthwhile.

The principal conclusion from this work is that the additional substitutions that have been permitted with the new **SeqStruct** substitution matrix enable obtaining a significantly stronger agreement between structure and sequence similarities. One of our current efforts is to establish a significantly more reliable substitution matrix beginning with the work reported here, but ensuring that the sequence matches doe not include significant false positive results.

The outcome of this research can improve the results in a large number of applications of sequence matching in biology, advancing particularly the fields of molecular, structural and evolutionary biology. A few of the many important gains will be improved gene annotations, vimproved protein structure determination, improved protein modeling, improved evaluations of the effects of protein mutants, and enhanced abilities to carry out reliable protein design. It is through this effort that the vast data from genome sequencing will come into practical application. Some problems that the present results may help to resolve:

1. Inability to recognize proteins having similar structures but dissimilar sequences
2. Mistakes in function identification based on previous sequence matches
3. Sequence matches that are not sufficiently informative about the effects of mutations

Many parts of biology depend on results from sequence matching, but these do not account for important structural properties such as amino acid packing, which provides the framework for the present work.

### Other ways to obtain gains

Manual inspection of the MSAs from Pfam shows that them not to be entirely reliable, and the MSAs themselves could be significantly improved by using the present **SeqStruct** similarity matrix. To obtain a better understanding of the functional mechanisms of a protein structure, it is important to integrate all available information to extract the complex co-evolution signals. Most previous studies have limited themselves to only pairwise correlations, but the present approach could be extended to consider larger groups - triplets, etc. It may be important to consider such higher order (higher than pairs) co-evolving dependencies because of the high packing density in proteins. We plan to do this in the future. Other significantly correlated clusters that are not proximate could also have been included, perhaps by including a distance weighting scheme. This would bring protein allostery directly into sequence matching. Allosteric spines that had been identified separately could even be weighted more strongly ^79-82^. Another way to make gains is to develop separate similarity metrics for different classes of protein folds. We have demonstrated this in a recent paper ^37^, but the gains appear to be modest.

### Huge Pent-up Needs

There are many specific applications that can benefit from improved sequence matching. For example, there are proteins involved in nuclear organization and cell division, where researchers have been unable to identify critical proteins these to any proteins in other organisms ^83^. Other scientists obtain crystals of a new protein but are unable to find a homologous known structure to aid in initial construction of a protein model. Comparative genomics across diverse species are often unable to identify the function of even half of the proteins. Applications of site-directed mutagenesis will likewise benefit from considering pairs of substitutions according to Table S2. These needs are only a small sample of the large number of research topics that can benefit from improved protein sequence matching.

### Potential Impacts

In summary, the approach applied here combines two extremely different sets of data – the protein sequences and structures, and combines them within a physical context by using close contacts in the structures to select the strongest correlations. This yields significant gains in sequence matching. The results produce dramatic improvements in sequence matching that will aid the present users of BLAST or any other present sequence matching software, because it relies on a simple 20×20 amino acid similarity matrix just as most other matching procedures also use. So its use is straightforward to implement. Further possible extensions include incorporating higher order multi-body correlations manifested in structures, may provide additional gains in the representations of the complex, dense proteins.

In general, biology is comprised of dense systems, with huge numbers of components, so the approach taken here may be generalizable to treat a wide range of specific complex problems. The critical data for applying such an approach would be the identities of the species and which ones are interacting with one another.

### The Future

There is a important need for efficient protein sequencing that would follow upon the successes with DNA and RNA sequencing. There is also the possible emergence of high throughput protein structure determination with free-electron X-ray crystallography. But despite this, there is likely to remain a substantial reliance on protein sequence matching.

## ACKNOWLEDGEMENT

This work was supported by the Carver Trust Grant to the Department of Biochemistry, Biophysics, and Molecular Biology.

